# CBAM-ResNet34-based classification and evaluation method for developmental processes of greenhouse strawberries

**DOI:** 10.1101/2024.12.03.626693

**Authors:** Jianxu Wang, Zhongyue Liang, Fengan Jiang, Jian Feng, Yuyang Xiao, Ming Yang, Deguang Wang

## Abstract

Strawberries, known for their economic significance and rich nutritional value, are cultivated extensively worldwide. However, a host of workers need to be employed every year to identify and categorize the developmental stages of the strawberries in the greenhouses, which is not only time-consuming, inefficient, increasing the cultivation cost, but also difficult to guarantee the classification accuracy. Meanwhile, affected by the complicated background, occlusions, and color interference, the features of strawberries are proven challenging to be extracted via the traditional neural networks due to serious gradient disappearance. Therefore, an improved CBAM-ResNet34- based classification evaluation method for developmental processes of greenhouse strawberries is investigated. The procedure of this method is as follows: firstly, the developmental stages of greenhouse strawberries are classified by experts into four stages: Stage I (initial stage), Stage II (green and white fruit stage), Stage III (early ripening stage), and Stage IV (fully ripe stage). The 627, 640, 604, and 340 strawberry images for these four stages are captured. Subsequently, the images are divided into training, validation, as well as testing sets and then undergo image pre- processing, expansion, and augmentation. Whereafter, the 7×7 convolution kernel in the first layer of the network is replaced by three consecutive 3×3 convolution cores to eliminate the redundant weights and unnecessary model parameters, and the BasicBlocks configuration is adjusted. Finally, the CBAM attention mechanism is added to each BasicBlock so as to pinpoint the spatial position of the strawberries and extract their major features such as shape, size, and color. Comparison experiments with the conventional deep neural networks LeNet5, AlexNet, VGG16, ResNet18, ResNet34, and every improved part of CBAM-ResNet34 demonstrated that when the learning rate is 0.001, the Dropout rate is 0.3, and the Adam’s weight decay parameter is 0.001, the accuracies for validation and testing sets can reach to 92.36% and 87.56% with F1 scores of 0.92, 0.87, 0.85 and 0.88.

## 1. Introduction

Strawberries, as a ubiquitous fruit abundant in nutritional value, are renowned for their richness in free sugars and organic acids (*Ikegaya*, 2023), rendering them suitable for the creation of a variety of delectable sweet and tangy products (*Oliveria* et al., 2021). Furthermore, they are recognized for their significant health benefits due to their diverse array of health-promoting active ingredients (*Tulipani* et al., 2009; *Newerli-Guz* et al., 2023), as well as their potential in anti-inflammatory, reducing the risk of heart and obesity-related ailments. (*Van de Velde* et al., 2019; *Gasparrini* et al., 2017; *Afrin* et al., 2016). Although the strawberries possess numerous merits mentioned above, a large number of strawberries are cultivated in warm and sunny greenhouses due to their demanding growing conditions. Therefore, during the strawberry growing stage and picking season in particular, a large number of workers are required to monitor and categorize the numerous and widely distributed strawberries in the greenhouse in real time (*Ibba* et al., 2021). Therefore, a fast, efficient, low-cost and objective method for the classification and evaluation of developmental stages of greenhouse strawberries is urgently required.

The ripeness of strawberries is intricately linked to their shape, size, and color (*Sturm*, et al., 2003). Therefore, pinpointing and highlighting their shape, size and color is the key to determine their maturity. The machine vision, as an advanced image processing and pattern recognition technology, has presented great potential and broad application prospects in various fields. Typical deep neural networks, such as AlexNet (*Krizhevsky*, et al., 2012), VGG (*Simonyan* & *Zisserman*, 2015), ResNet (*He*, et al., 2016), DenseNet (*Huang*, et al., 2017), and Inception V4 (*Szegedy*, et al., 2017), have been widely applied and popularized in agricultural fields such as fruits and vegetables. Not only can a great variety of food types and quality levels be classified by these conventional deep neural networks (*Zhang* et al., 2014), but also their nutritional content and developmental processes can be determined (*De* et al., 2022; *Ma* et al., 2021; *Schieck* et al., 2023), and pests and diseases in crops can be monitored and prevented (*Li* et al., 2020; *Mahmud* et al., 2020; *Guo* et al., 2022; *Too* et al., 2019). Deep neural networks, however, are plagued by the issue of vanishing gradients with an increasing number of layers, and are prone to being perturbed by factors such as variable backgrounds, occlusion by foreign objects, and interference from colors, leading to compromised performance in the classification and evaluation of strawberry ripeness. Therefore, an efficient network that is not only in a position to extract pivotal features of strawberry shape, size, and color, but also capable to effectively learning deeply is urgently needed.

Currently, notable classification results have been attained in certain studies through the utilization of machine learning methods for feature extraction in image color space, with key features combined with strawberry ripening stages into a dataset for training (*Karki* et al., 2023; *Wise* et al., 2022). Similarly, the features such as diameter, length and top angle of strawberries were extracted by image processing algorithms, and were then employed as classification indicators for model training. The estimation of strawberry shape and size was thus effectively enabled through this approach (Oo & Aung, 2018). Nevertheless, it leads to additional challenges of intricate and limited feature extraction, lengthy time consumption, and difficulty for handling environmental background interference, which yet remains unresolved.

Since color being a major indicator determining food quality, a comparative analysis of the food color distribution via the machine vision method and sensory perception was conducted (*Ayustaningwarno*, et al., 2021), and the results indicated that machine vision outperformed manual judgement, and its implementation is not only characterized by flexibility and non-destructiveness, but also by greater cost-effectiveness and economic viability. In addition, the volume estimation of agricultural products with irregular shapes has been materialized based on machine vision method (*Concha-Meyer*, et al., 2018). Furthermore, Mask R-CNN and region segmentation techniques were effectively utilized in the task of strawberry ripening classification with environmental occlusion. (*Tang* et al., 2023). Similarly, the combination of Faster R-CNN with the CBAM attention mechanism was employed to effectively localize and extract the position and disease characteristic information of strawberries, and it greatly enhanced the management efficiency of farmers during the cultivation of strawberries (*Zhao* et al., 2022).

To address the problems of time-consuming, inefficient, costly and subjective manual evaluation of the developmental stages of greenhouse strawberries, the poor performance of deep neural networks for classification tasks with high environmental disturbances (e.g., occlusion, light imbalance and background similarity), and the disappearance of gradient due to deepening of the layers, a CBAM-ResNet34-based classification evaluation method for developmental processes of greenhouse strawberries is investigated. Firstly, the edge shape and surface detail features of strawberries were highlighted by sharpening (*Fang* et al., 2008), histogram equalization, and other operations, and the dataset was expanded by cropping, flipping, and rotating operations. Whereafter, the residual connection of the ResNet34 is adopted to solve the problem of gradient vanishing, and the CBAM attention mechanism is added to each BasicBlock to accurately locate the position of the strawberries and highlight their shape (*Woo* et al., 2018), size and color information through spatial and channel enhancement. Subsequently, the 7×7 convolution kernel in the first layer of the network is replaced by three consecutive 3×3 convolution cores to extract deep features, and the number of BasicBlock is fine-tuned to be 3, 4, 2, and 2 (*Li* et al., 2022), retaining the shallow as well as critical feature extraction layers to reduce the complexity, which can achieve the balance between computation and accuracy.

The research contributions of this study are as follows:

1. The time-consuming, inefficient, costly and subjective problems of manually experience judging strawberry maturity can be solved by the artificial intelligence technologies.
2. The gradient disappearance that existed in the traditional deep neural networks with the increase of layers is effectively solved by utilizing the residual structure. Meanwhile, the key feature information such as shape, size and color of strawberries are highlighted by the image sharpening and other enhancement operations.
3. The position of the strawberry is successfully localized by introducing CBAM attention mechanism, and then the key feature information is deeply mined. Additionally, the environmental background interference is filtered out, which further improves the classification accuracy.

## 2. Materials and Methods

### 2.1 Strawberry sample image dataset acquisition

Strawberry samples were obtained from strawberry greenhouses in Donggang, Dandong city, Liaoning Province, China, and the images collection sketch is depicted in Figure 1.

**Figure 1.**
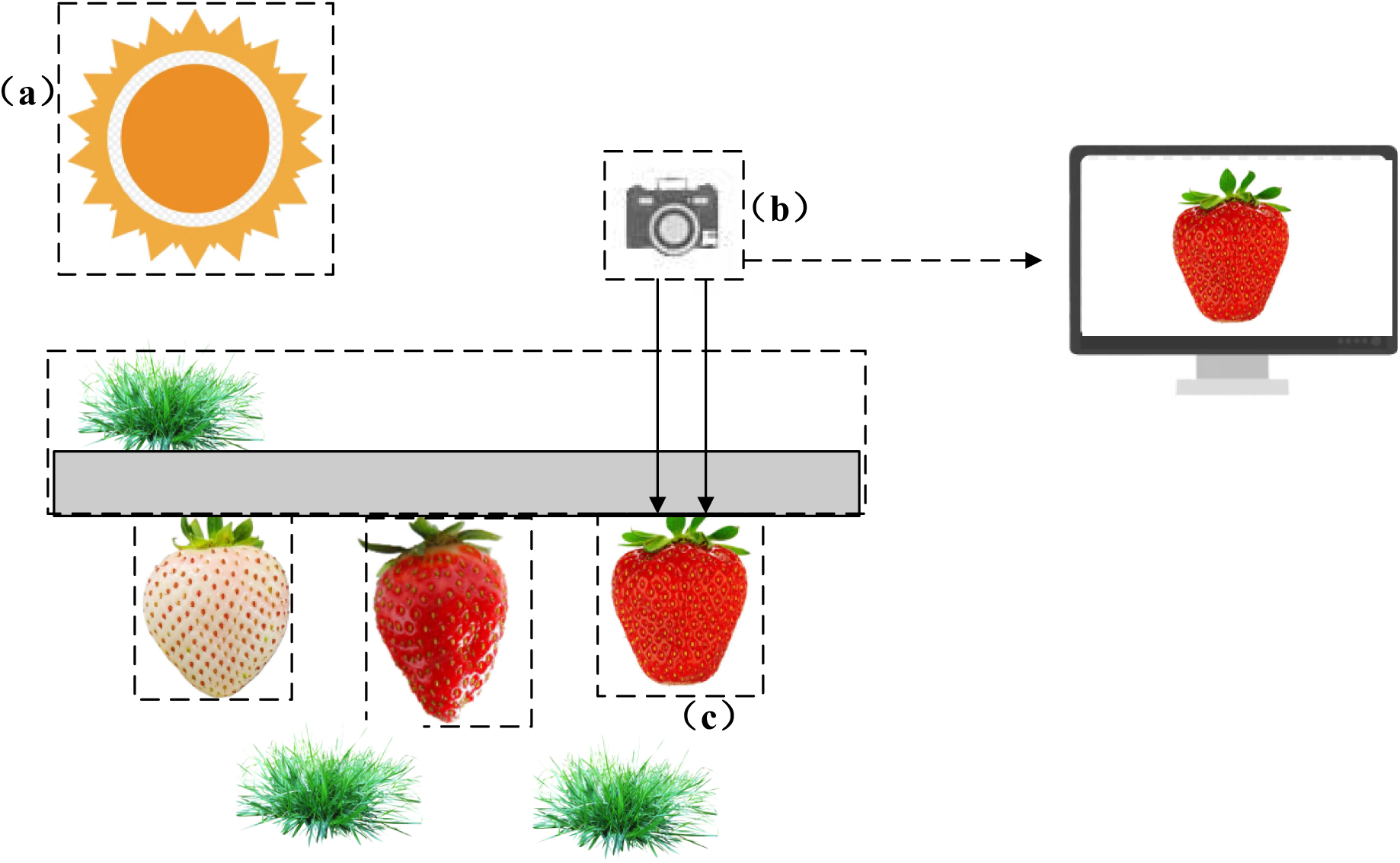
Sketch of strawberry sample image data acquisition scene.

The image collection processing is as follows: firstly, the experts entered the strawberry greenhouses to evaluate the developmental process of strawberries and classify them into four stages: Stage I (initial stage), Stage II (green and white fruit stage), Stage III (early ripening stage), and Stage IV (fully ripe stage). Secondly, the light (a) in the shed was appropriately adjusted during the collection process to emphasize the surface characteristics of strawberries. Subsequently, in order to enhance the model’s ability of resisting interference with the background and environment, the shooting distance between the camera (b) and the strawberry (c) and the shooting angle are adjusted. Finally, the 2211 strawberry images of four developmental stages were obtained, and they were sequentially divided into training set, validation set and test set. The four classes of strawberry sample datasets were divided, as listed in Table 1, and the schematic diagram of the four classes of strawberry developmental stage images was depicted in Figure 2.

**Figure 2.**
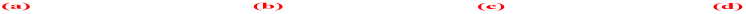
Diagram of four types of strawberry maturity: (a) Stage I, (b) Stage II, (c) Stage III, (d) Stage IV.

**Table 1.** Distribution of training set, validation set, and testing set.

| Developmental stage | Total number | Training set | Verification set | Testing set |
| --- | --- | --- | --- | --- |
| Stage I | 340 | 200 | 70 | 70 |
| Stage II | 604 | 444 | 80 | 80 |
| Stage III | 640 | 320 | 160 | 160 |
| Stage IV | 627 | 347 | 140 | 140 |

### 2.2 Image preprocessing

When obtaining strawberry images, the 1 to 1 shooting was adopted to prevent the shape from being compressed when scaling the image and causing the model to lose its learning ability, and the image pre-processing operation is depicted in Figure 3, which operates as follows: firstly, the sample images of strawberries are uniformly scaled to 255×255 based on the Pytorch function. Secondly, the scaled images are randomly flipped horizontally and randomly flipped vertically to expand the training set. Then, the images are further transformed into computer-processable Tensor form. Finally, in order to avoid a large integer value slowing down the model training speed, these tensor-type data will be further normalized by:

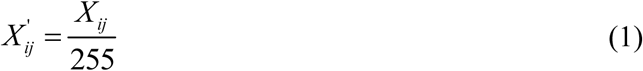

**Figure 3.**
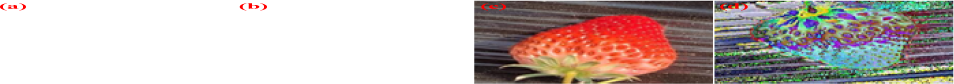
Image pre-processing operation: (a) original image, (b) random clipping, (c) flip horizontal, (d) normalization and standardization.

where, *X_ij_* (*i* = 1, 2, 3; *j* = 1, 2, …, 224x224) is the brightness value of the *i*th channel of the *j*th pixel point in the original image. These data will then be further standardized by:

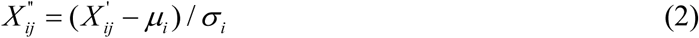

of which, *μ_i_* is the brightness average value of the three channels, which is 0.485, 0.456, and 0.406. *σ_i_* is the standard deviation of the brightness values of the three channels, which is 0.224, 0.229, and 0.225, respectively. Each image preprocessing operation is displayed in Figure 3.

## 3. Construction of CBAM-ResNet34

### 3.1 Traditional ResNet

Remarkable success has been achieved in the fields of image recognition and speech recognition through the application of deep residual networks (*He* et al., 2016). Consequently, ResNet34, known for its efficiency with reduced computational and model parameters that effectively tackle the gradient vanishing through a residual series, was chosen as the foundational model for this study. The BasicBlock structure of ResNet34 is illustrated in Figure 4, which constructs the residual function *F*(*x*) from two 3×3 convolution kernels, and is designed by the following simple mapping before input samples *x*:

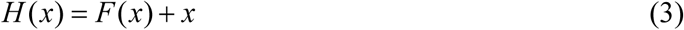

**Figure 4.**
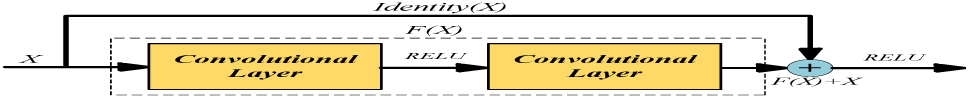
Schematic diagram of the BasicBlock.

when *F(x)* = 0, an identity map *H(x) = x* is formed. The network layer is also an identity mapping, the network performance doesn’t degrade, and the residual fitting become easier.

### 3.2 CBAM attention mechanism

The CBAM is an attention mechanism commonly applied in image classification (*Woo*, et al., 2018). It is constituted by a channel attention mechanism module (CAM) and a spatial attention mechanism module (SAM), as depicted in Figure 5. In the feature extraction stage of strawberries, the shallow feature maps with rich semantic information are utilized to extract macroscopic features such as shape, texture, and contour of strawberries, while the deeper feature maps are employed to capture abstract semantic information. However, the detailed features such as the shape and size texture of strawberries are easily neglected after multiple deep convolutions, and the corresponding gradient information is thus thrown away. To solve the problems mentioned above, the CBAM attention mechanism is introduced to carry out attention weighting and information gathering. Where, the channel attention *A_c_* can be expressed as:

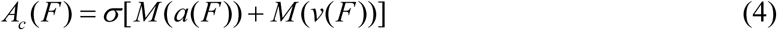

**Figure 5.**
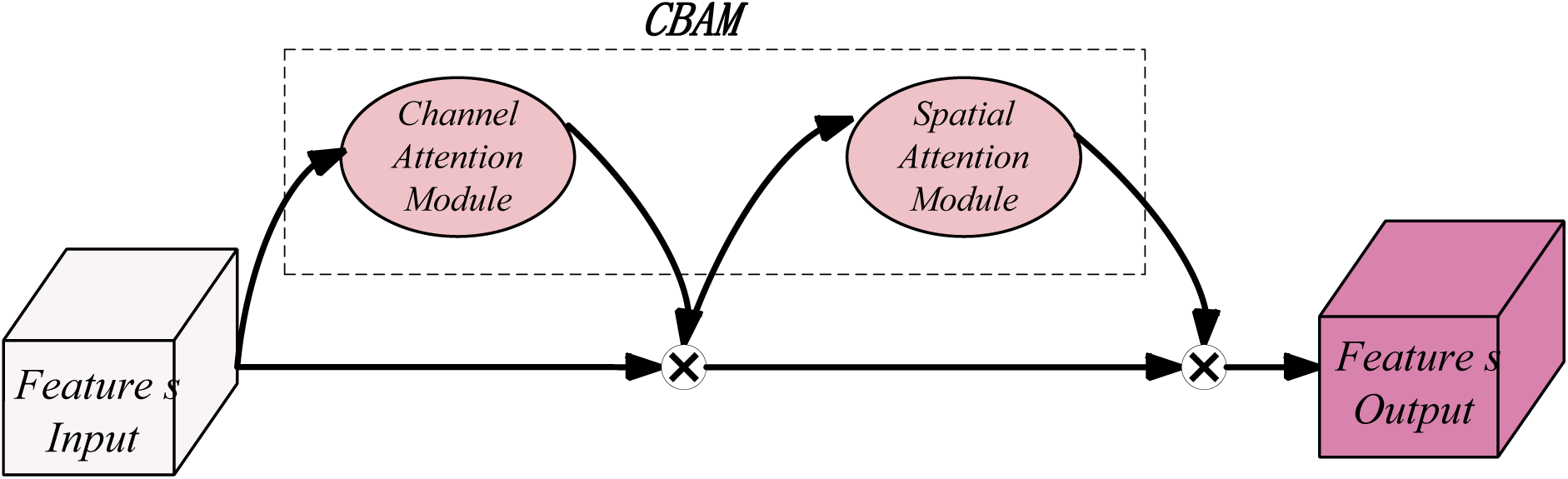
Schematic diagram of the CBAM.

where, *F* is the input feature map, a and v are the maximum and average pooling operations, *M* is the multi-layer perceptron operation, and *σ* is the Sigmoid operation. Where, the spatial attention is calculated by:

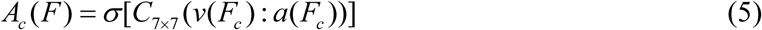

of which, *F_c_*is the feature map obtained by CAM, *C_7×7_* is the convolution operation with a kernel size of 7x7.

Within the attention mechanism module, the CAM is employed to reinforce the learning of intricate features, such as shape, texture, degree of gloss, color, and size of strawberries, while the SAM enhances discrimination between strawberries, background, and occluded regions. The attention map resulting from their fusion is subsequently multiplied by the original feature map to derive the comprehensive weight map highlighting the focal areas. They are calculated by:

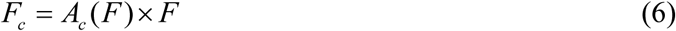

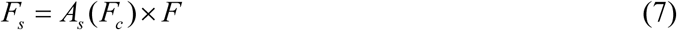

where, *F_s_*is the corresponding attention map.

### 3.3 Fine-tuning of initial convolution and BasicBlock layers

The extraction of detailed features of strawberry has an important impact on the taxonomic evaluation of developmental processes. Nevertheless, the conventional ResNet34 model’s initial input layer utilizes a 7×7 convolutional kernel that retrieves extensive neighborhood range information from the entire strawberry image, which is unfavorable for detailed feature extraction. Therefore, three consecutive 3×3 convolutional kernels are employed in place of the 7×7 convolutional kernels in the initial layer, and the illustration comparing the convolution layers before and after the enhancement is depicted in Figure 6.

**Figure 6.**
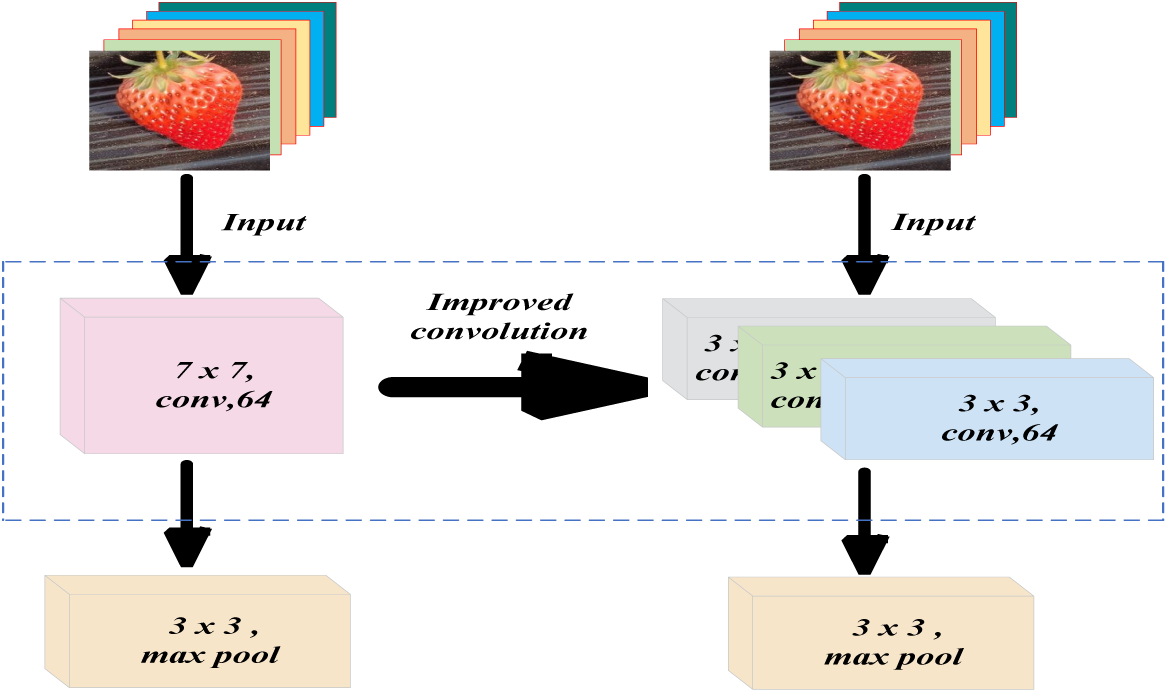
Improvement of initial convolution.

The sensory field is the size of the region where pixels on the feature map are mapped on the input image at each output during the convolution process, which is calculated as:

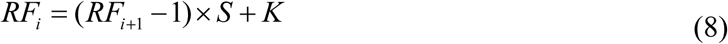

According to the computational results, the improved three 3×3 convolutional kernels have the same receptive fields as the original 7×7 convolutional kernels. However, the use of three 3×3 convolutional kernels increases the depth of the network, and with the same number of input channels, the ratio of the number of parameters in the model before and after the improvement is (7×7) : (3×3×3) = 49 : 27, and the number of parameters in the pre-improvement period is only about 55% of the number of parameters in the post-improvement period, which improves the speed of operation of the network.

In comparison with the traditional ResNet34, this paper fine-tunes the number of BasicBlocks in each layer to 3, 4, 2, and 2, respectively, with the aim of eliminating redundant features and network weights, significantly compressing space, and reducing the number of parameters and computations (*Li* et al., 2022).

### 3.4 Improved ResNet34

Firstly, the image preprocessing, image enhancement and image expansion are utilized, the diversity of strawberry samples is enhanced, and features related to ripeness such as size, shape and color of strawberries are highlighted. Secondly, three consecutive 3×3 convolutional kernels are utilized instead of the 7×7 convolutional kernels of the initial layer, where the first convolutional kernel with a step size of 2 is used to generate an intermediate image of 112×112×64 by extracting the features from the input image of 224×224×3, while the second and the third convolutional kernels with a step size of 1 are used to keep the image size of 112×112×64. Subsequently, the batch normalizeation is performed (*Ioffe* & *Szegedy*, 2015) and the ReLU activation is applied, and the image features were further extracted by a 3×3 convolution kernel, followed by a maximum pooling operation with a step size of 2 and a zero padding of 1 to generate a 56×56×64 scaled image. Then, the image was sequentially passed through 3, 4, 2, and 2 layers of improved BasicBlock residual blocks (*Li* et al., 2022), and the CBAM attention mechanism (*Woo* et al., 2018) was added before the BasicBlock of each Layer to enhance the expression of features. With these operations, a final 7×7×512 image was generated.

Then, the image was sequentially passed through 3, 4, 2, and 2 layers of improved BasicBlock residual blocks (*Li* et al., 2022), and the CBAM attention mechanism (*Woo* et al., 2018) was added before the BasicBlock of each Layer to enhance the expression of features. With these operations, a final 7×7×512 image was generated. Finally, the image is scaled to 1×1× 12 using an adaptive average pooling layer and a newly designed three-layer fully connected layer with Dropout is introduced (*Srivastava* et al., 2014) to better extract the feature information of the strawberries and improve the recognition accuracy and prevent network overfitting. The features are further extracted in the last two layers, and the optimal deactivation rate is selected by setting the proportion of randomly deactivated nodes in Dropout, and the final classification results are output. The schematic diagram of the overall model planar architecture is shown in Figure 7, and the schematic diagram of the three-dimensional architecture is displayed in Figure 8.

**Figure 7.**
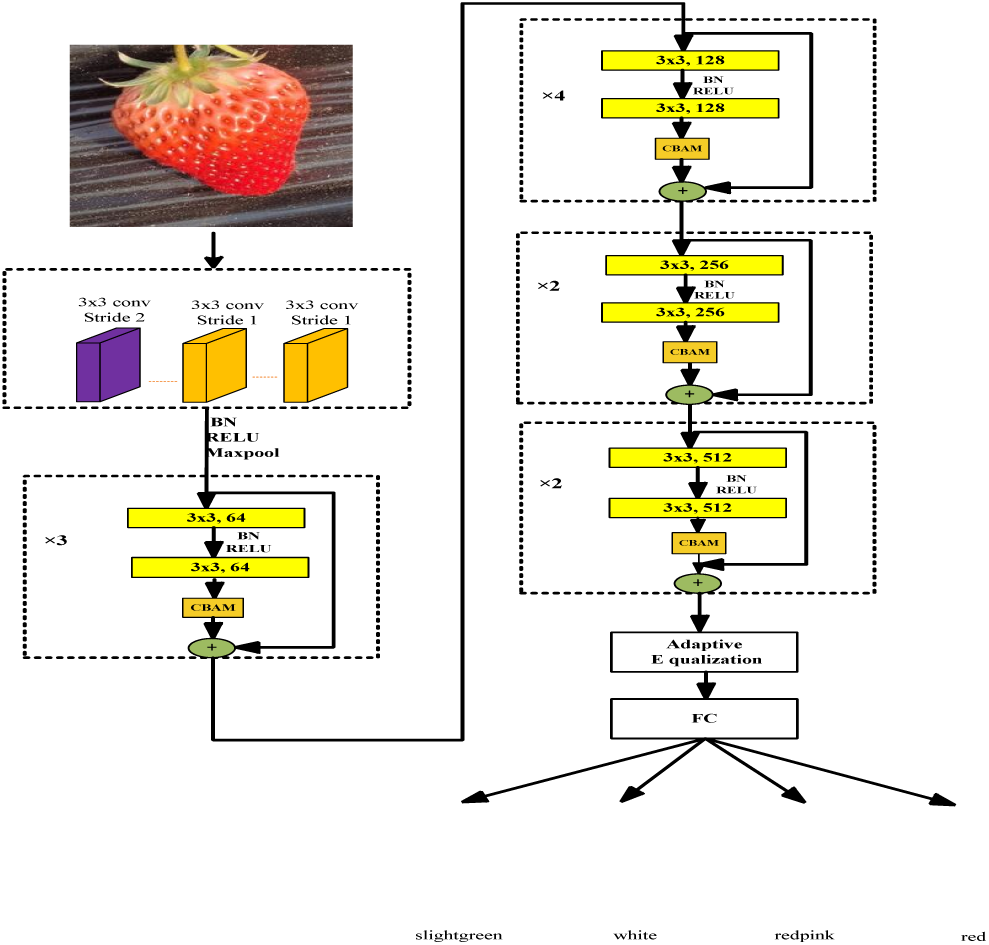
Plane architecture of the CBAM-ResNet34 network.

**Figure 8.**
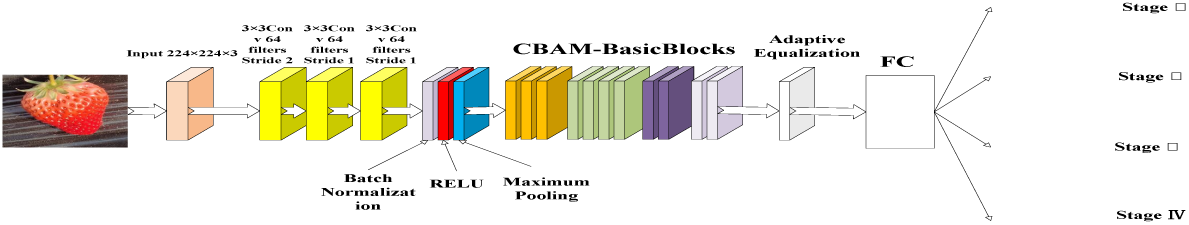
Solid architecture of the CBAM-ResNet34 network.

## 4. Experiments and Results

### 4.1 Experimental setup

The experimental conditions were: NVIDIA GeForce RTX 3050 Laptop GPU with 11071 MB RAM, 3977 MB video memory, 7094 MB shared memory, Windows operating system. The Pytorch framework was used to build and train deep learning models, and the adopted version and cuda of torch, torchaudio, and torchvision versions were 1.12.0+cu113, 0.12.0+cu113, and 0.13.0+ cu113, respectively. The code was programmed on the software PyCharm.

### 4.2 Performance comparison of improved model

4.2.1 Comparison between the improved model and traditional network models In order to verify the comprehensive performance, five classical networks were in comparison with the CBAM-ResNet34: LeNet5, AlexNet, VGG16, ResNet18, and traditional ResNet34. The training times were set to 200, with a learning rate of 0.001, Dropout rate of 0.5, Adam optimizer, and cross-entropy loss function. And the experimental outcomes are provided in Table 2.

**Table 2.** Performance comparison of different models.

| Model | Number of parameters | FLOPS | Accuracy of verification set | Accuracy of testing set | Total training time |
| --- | --- | --- | --- | --- | --- |
| LeNet5 | 61720 | 0.0023G | 86.28% | 78.89% | 6min11s |
| AlexNet | 61.10M | 0.71G | 87.54% | 79.78% | 22min23s |
| VGG16 | 138.36M | 15.47G | 88.63% | 80.67% | 14h49min |
| ResNet18 | 11.69M | 1.82G | 88.87% | 81.56% | 25min55s |
| ResNet34 | 21.80M | 3.68G | 89.68% | 82.67% | 40min48s |
| CBAM-ResNet34 | 23.66M | 3.81G | 92.36% | 87.56% | 49min43s |

Affected by the challenges of intricate environmental interference of the shooting scene (random background, occlusion, and similar color), the traditional LeNet5, AlexNet, and VGG16 show limited improvement in accuracy rates. However, ResNet18 and ResNet34 achieved an accuracy of over 81% through effective residual link. Furthermore, by incorporating the CBAM attention mechanism, the model effectively eliminated environmental interference, identified the strawberry’s location, and extracted its appearance features, leading to an accuracy improvement to 86.22%. Finally, through parameter tuning based on model characteristics, CBAM-ResNet34 attains an inference accuracy of 87.56%, which demonstrates that CBAM-ResNet34 has a notable advantage in coping with complex background interference.

The training loss and validation set accuracy of the seven models are depicted in Figures 9 and 10, respectively. Due to their shallower network hierarchies, both LeNet5 and ResNet18 exhibit faster convergence speeds and lower final convergence losses compared to the ResNet34, AlexNet, and VGG16, especially after fine-tuning BasicBlock in CBAM-ResNet34. While LeNet5 falls short in achieving higher validation set accuracy due to its network depth and limitations, CBAM-ResNet34 leverages the residual network to enhance model generalization and inference accuracy effectively. Most importantly, with the integration of CBAM’s attention mechanism, it excels in pinpointing the strawberry’s location in new scenarios and capturing its appearance features, resulting in a nearly four percentage point increase in test set accuracy.

**Figure 9.**
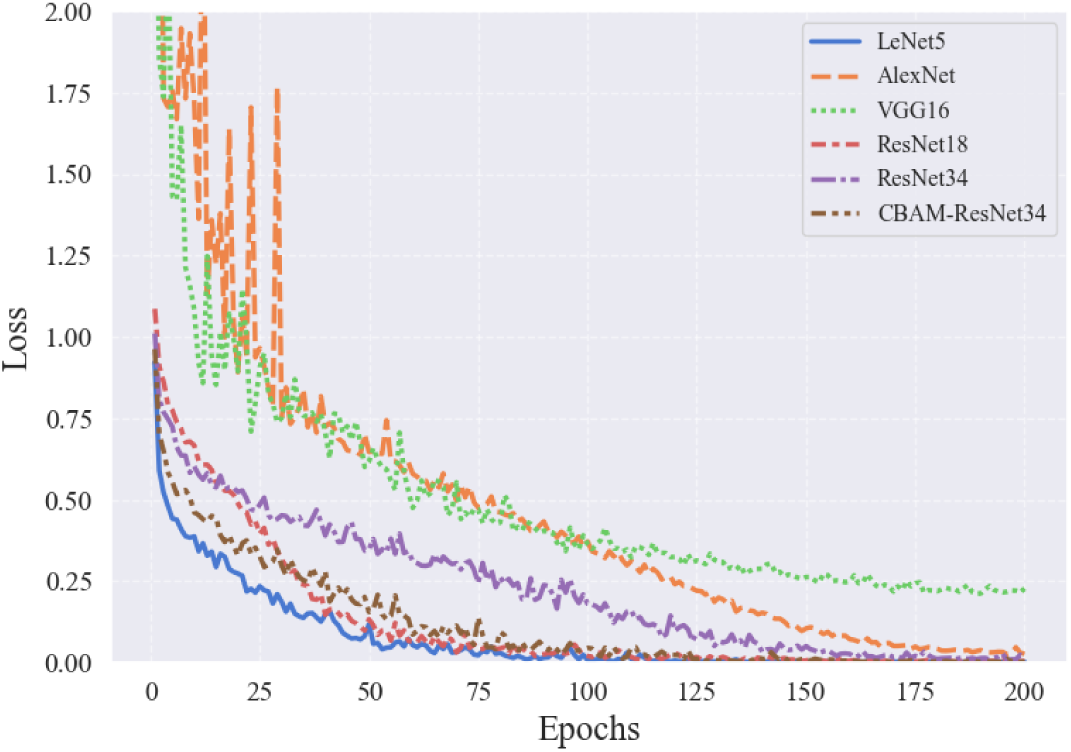
Loss function of different network models.

**Figure 10.**
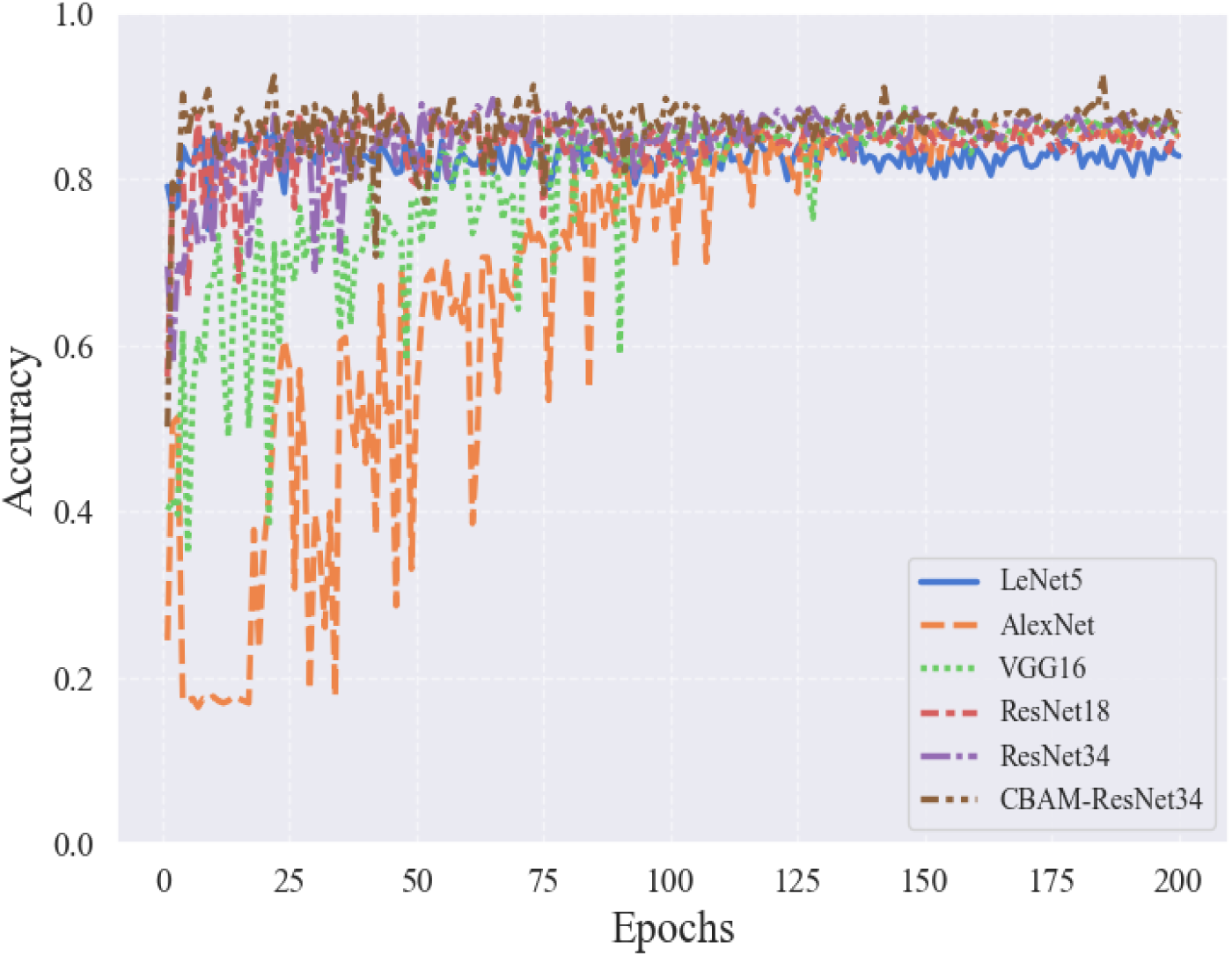
Verification set accuracy of different network model.

In contrast, the deeper architectures of AlexNet and VGG16, with more parameters and sensitivity to initial weights, exhibit fluctuating training loss and validation curves, necessitating longer training times for stable convergence. In summary, the fine-tuned ResNet34 with CBAM stands out from traditional models, showcasing superior classification performance in assessing strawberry maturity.

4.2.2 Ablation experiment

To elucidate the impact of incorporating the CBAM-based fine-tuning method on ResNet34, the base model ResNet34 is augmented by adding the CBAM attention mechanism separately and enhancing the initial convolution. The combined fine-tuned network is then compared with these variations. Experimental results are provided in Table 3 for evaluation.

**Table 3.** Ablation experimental results.

| Model | Accuracy for verification set | Accuracy for testing set |
| --- | --- | --- |
| Traditional ResNet34 | 89.68% | 82.67% |
| CBAM | 89.58% | 84.22% |
| Improved initial convolution | 89.84% | 82.89% |
| [3, 4, 2, 2] | 90.10% | 82.89% |
| CBAM + [3, 4, 2, 2] | 91.41% | 85.11% |
| CBAM + Improved initial convolution | 90.36% | 84.44% |
| [3, 4, 2, 2] + Improved initial convolution | 88.54% | 83.33% |
| Improved ResNet34 | 92.62% | 86.22% |

In analyzing the test set performance results, it is evident that the CBAM attention mechanism plays a crucial role in significantly extracting the position and surface features of the strawberry, thereby greatly enhancing the model’s generalization ability. On the other hand, the enhancements in the initial convolution and fine-tuning of BasicBlock layers primarily contribute to improving model performance, to some extent reducing model complexity and mitigating the risk of overfitting.

### 4.3 Impact of Different Learning Rates, Dropout and Adam Weight Decay

In this experiment, five different sets of learning rates: 0.01, 0.001, 0.005, 0.0001, and 0.00001 are set up to investigate the effect of learning rate on model performance (*Liu* et al., 2022). The experimental results were listed in Table 4: learning rates of 0.01, 0.005, and 0.001. Nevertheless, as the learning rate decreases further, the model’s performance deteriorates, leading to convergence to local optima and significantly compromising its generalization capability. Hence, a learning rate of 0.001 is deemed the optimal parameter choice.

**Table 4.** Validation, test set accuracy of improved models at different learning rates.

| Learning rate | Accuracy for verification set | Accuracy for testing set |
| --- | --- | --- |
| 0.01 | 90.32% | 85.56% |
| 0.001 | 92.62% | 86.22% |
| 0.005 | 90.89% | 82.00% |
| 0.0001 | 90.36% | 83.33% |
| 0.00001 | 87.76% | 81.56% |

Whereafter, five different Dropout inactivation rates are set: 0.3, 0.4, 0.5, 0.6, and 0.7 (*Srivastava* et al., 2014), and the results of the experiment were provided in Table 5.

**Table 5.** Validation, test set accuracy of improved models at different Dropout rates.

| Dropout rate | Accuracy for verification set | Accuracy for testing set |
| --- | --- | --- |
| 0.3 | 91.32% | 86.67% |
| 0.4 | 89.32% | 84.00% |
| 0.5 | 92.62% | 86.22% |
| 0.6 | 90.63% | 83.56% |
| 0.7 | 89.58% | 84.22% |

The experimental findings demonstrate that the model attains superior performance with Dropout inactivation rates of 0.3 and 0.5. However, the generalization capacity is weakened when the Dropout inactivation rate reaches 0.5 due to over-inhibited neurons. On the other hand, a Dropout inactivation rate of 0.3 leads to a notable enhancement of the model’s generalization performance, provided that overfitting is effectively suppressed. Consequently, the optimal parameter choice is a Dropout inactivation rate of 0.3.

Eventually, four different sets of weight decay parameters are designed based on the Adam optimizer: 0, 0.001, 0.0005, and 0.0001 (*Zhou* et al., 2023), and the experimental results were listed in Table 6.

**Table 6.**
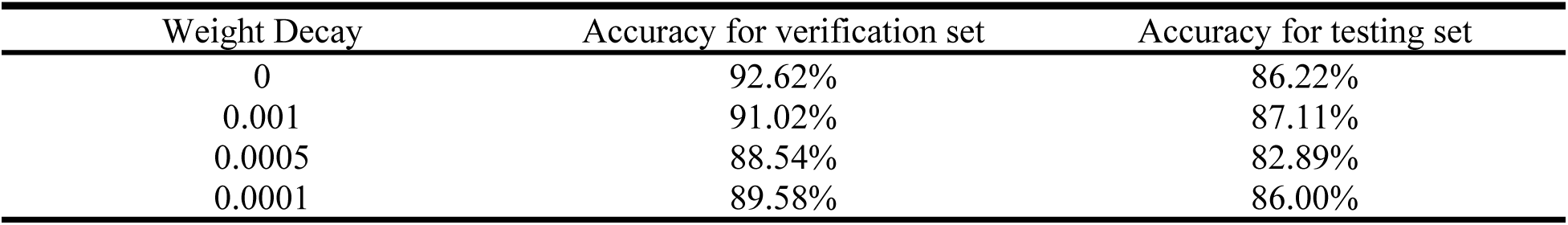
Improved model validation, test set accuracy under different weight attenuation based on Adam.

The experimental results reveal that with an increase in the Weight Decay value, the model’s loss gradually decreases at a slower pace. A Weight Decay value of 0.001 leads to a 0.89% improvement in the model’s performance on the test set, suggesting that the model complexity is constrained, overfitting is mitigated, and more robust and generalizable features are extracted to better align with real data distribution. Therefore, the Weight Decay of 0.001 is identified as the optimal parameter choice.

### 4.4 Overall performance of the improved model with optimal parameters

To investigate the combined effects of key parameters on the improved base model, optimal parameter experiments are conducted on the improved base model with the following parameters: 200 training rounds, 0.001 learning rate, 0.3 Dropout rate, and 0.001 Adam optimizer weight attenuation parameter. Eventually, a verification set accuracy of 92.36% and a testing set accuracy of 87.56% is obtained.

The experimental results indicate that with lower Dropout inactivation rates and larger Adam weight decay parameters, although the model’s losses decrease at a slower rate and exhibit more fluctuations in the validation set curves, there is a significant improvement in test set performance. This improvement can be largely attributed to the effective inhibition of the model’s overfitting through the regularization method of weight decay. Consequently, the model becomes more responsive to noise or changes in the validation set, enabling the extraction of more generalized and robust features that better align with the distribution of real data.

In summary, while the CBAM-ResNet34-based model with optimal parameters may not have shown a remarkable enhancement in validation set accuracy, it successfully extracts universal and robust features. The notable accuracy enhancement on the test set suggests a significant improvement in the model’s generalization capability.

### 4.5 Model performance assessment

In order to evaluate the effect of the CBAM-ResNet34-based classification evaluation method for developmental processes of greenhouse strawberries, four cases are utilized to showcase the classification results: true positive (TP), false positive (FP), true negative (TN), and false negative (FN). The precision (P), recall (R), and F1-score are utilized to evaluate the model’s classification performance for various developmental stages, and the accuracy, macro avg., and weighted avg. are adopted to evaluate the model’s overall performance. They are expressed as:

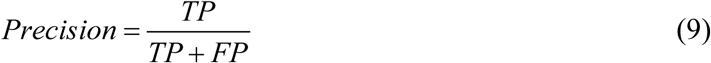

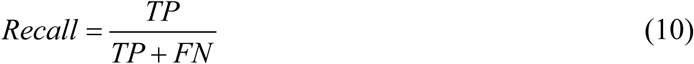

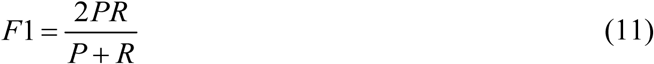

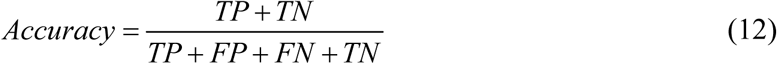

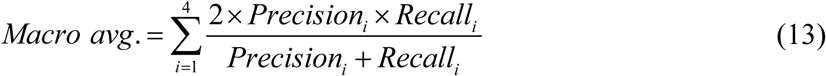

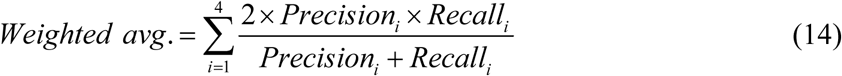

where, *N* represents the total number of samples, which is 450, and *N_i_* represents the number of Class *i*th samples.

The optimal parameter model based on CBAM-ResNet34 was predicted for the test sets with the total number of samples of 70, 80, 160, and 140 for stages I, II, III, and IV, respectively, and the heat maps of strawberry visualization for the four stages of growth and the corresponding inference results were obtained as illustrated in Figures 11 and 12, and the classification report was listed in Table 7.

**Figure 11.**
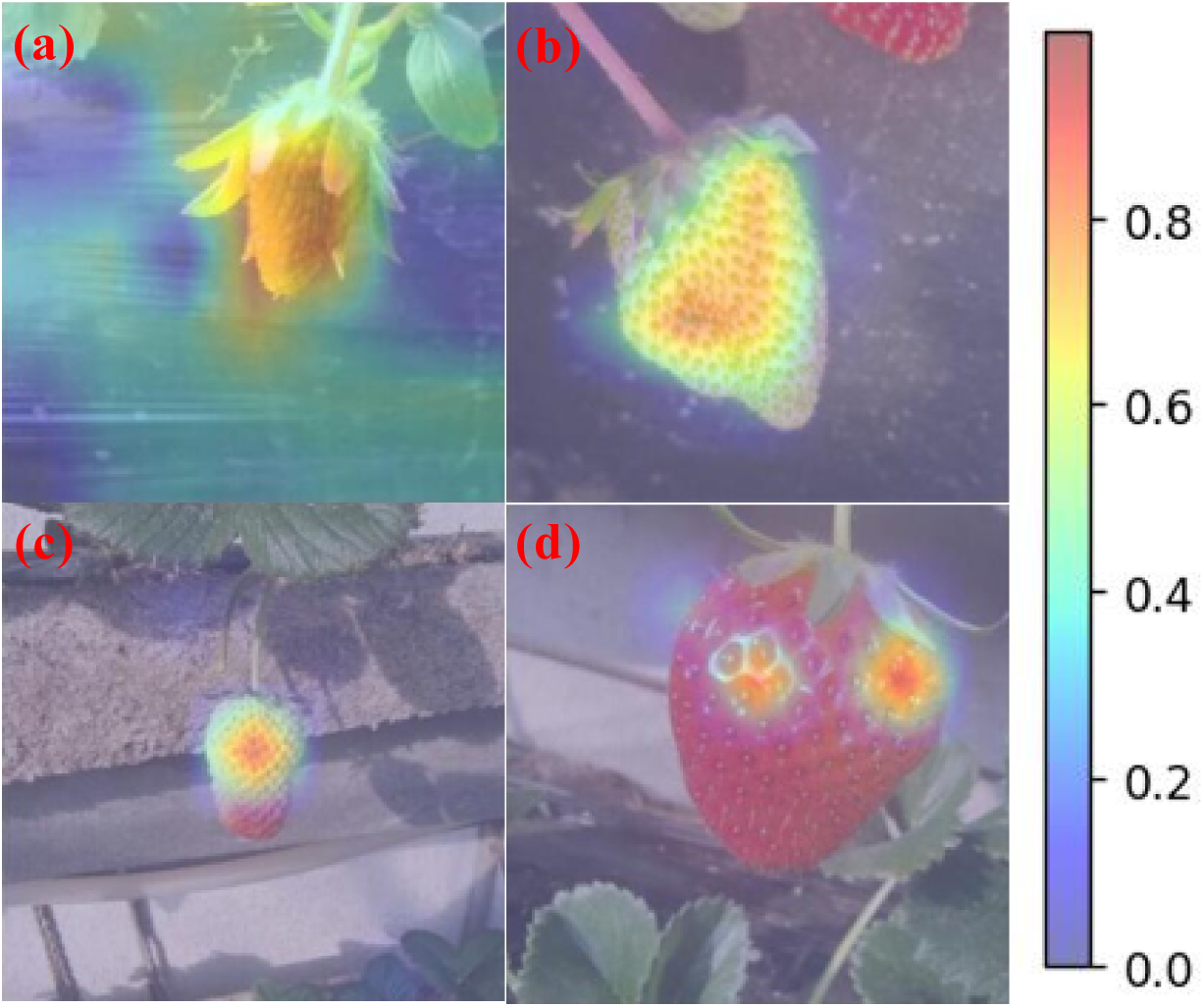
Visualized heat maps of four different developmental stages of strawberries: (a) Stage I, (b) Stage II, (c) Stage III, (d) Stage IV.

**Figure 12.**
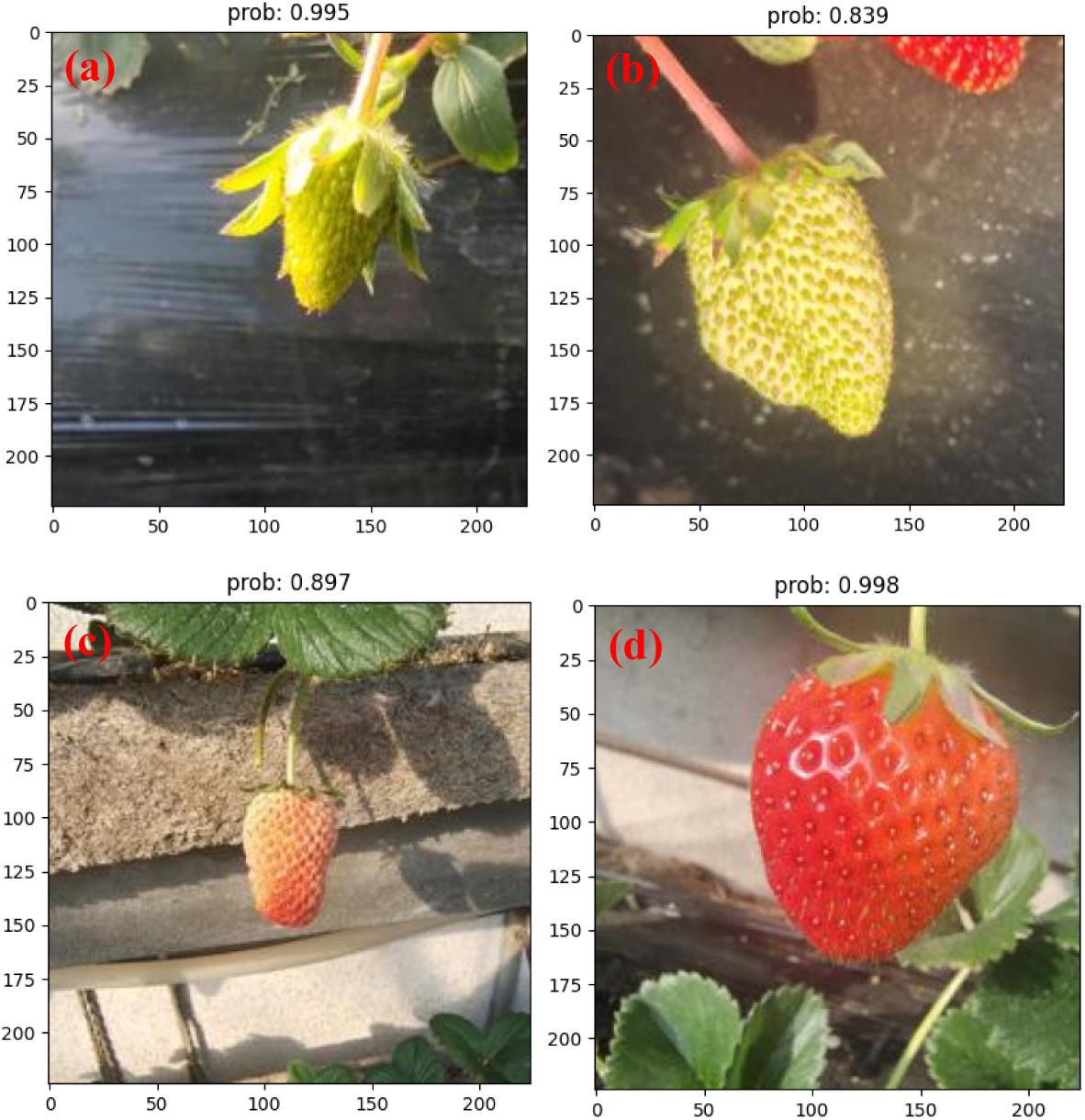
Model predictions for four developmental stages of strawberries: (a) Stage I, (b) Stage II, (c) Stage III, (d) Stage IV.

**Table 7.** The classification results by the CBAM-ResNet34 model.

| Category | Precision | Recall | F1-Score | Number of sample |
| --- | --- | --- | --- | --- |
| Stage I | 0.92 | 0.93 | 0.92 | 70 |
| Stage II | 0.86 | 0.88 | 0.87 | 80 |
| Stage III | 0.94 | 0.78 | 0.85 | 160 |
| Stage IV | 0.81 | 0.96 | 0.88 | 140 |
| Accuracy | / | / | 0.88 | 450 |
| Macro avg. | 0.88 | 0.89 | 0.88 | 450 |
| Weighted avg. | 0.88 | 0.88 | 0.87 | 450 |

Upon analysis of the aforementioned results, the precise localization of strawberries and comprehensive extraction of their key features are facilitated through the investigation of CBAM- ResNet34. Furthermore, as provided in Table 7, despite Stage III exhibiting a lower F1 score and posing relative classification challenges compared to Stage II and Stage IV, the Macro avg. of 0.88 and Weighted avg. of 0.87 indicate strong overall and weighted performance of the improved model. In conclusion, the enhanced model, achieved by integrating CBAM into traditional ResNet34 and fine-tuning the network, showcases exceptional classification capabilities, robust generalization, and resilience in classifying strawberry growth stages.

## 5 Conclusion

To solve the current challenges of time-consuming, inefficient, costly, and subjective manual classification and evaluation of greenhouse strawberry development, a CBAM-ResNet34-based comprehensive and efficient method for assessing strawberry ripeness was proposed. This method incorporates the CBAM attention mechanism to enhance the extraction accuracy of strawberry position, shape, size, and color features, integrates three consecutive 3x3 convolution kernels in the initial layer to reduce the parameters and computational burden, and adjusts the number of BasicBlocks in the four-residual layers from 3, 4, 6, and 3 to 3, 4, 2, and 2 successively. Comparison with the traditional deep neural networks, the proposed method demonstrated an approximate 10% improvement in inference accuracy and significantly enhances generalization capabilities.

Comparative experiments involving different learning rates, Dropout deactivation rates, and weight decay parameters were conducted, revealing that the enhanced model has the optimal classification performance when utilizing a learning rate of 0.001, a Dropout deactivation rate of 0.3, and an Adam-based weight decay parameter of 0.001. Moreover, the proposed method for the validation and testing sets has accuracies of 92.36% and 87.56%, respectively. The F1 scores for strawberry classification across stages I, II, III, and IV are determined as 0.92, 0.87, 0.85, and 0.88, with Macro avg. achieving 0.88 and Weighted avg. reaching 0.87.

This proposed method not only addresses the current challenges of manual evaluation of greenhouse strawberry developmental stages, including time consumption, inefficiency, high costs, and subjectivity but also tackles the issues faced by deep neural networks in classifying tasks under high environmental disturbances and gradient vanishing with increased layer depth. These findings hold significance for the classification and evaluation of developmental processes in various agricultural domains. Importantly, our study demonstrates promising transferability for classifying and evaluating developmental processes in other agricultural fields. Future research directions may involve exploring more sophisticated and precise classification models and developing lightweight handheld application devices based on this method to enhance its practical application in real-world agricultural settings.

## Data availability

The datasets used and/or analyzed during the current study available from the corresponding author on reasonable request.

## Acknowledgements

This work was supported in part by National Natural Science Foundation of China (No. 52265066, 62203132, 62341303); Youth Science and Technology Talents Development Project of Guizhou Education Department (No. Qianjiaohe KY [2022]138); Doctor Foundation Project of Guizhou University (No. GuidaRenjiHezhi [2020]30).

## Author contributions

Jianxu Wang and Ming Yang conceived and designed the study and wrote the paper. Jianxu Wang, Ming Yang, Deguang Wang, Zhongyue Liang, Fengan Jiang, Jian Feng and Yuyang Xiao performed the experiments. Jianxu Wang and Ming Yang reviewed and edited the manuscript. All authors reviewed the manuscript.

## Additional information

Correspondence and requests for materials should be addressed to Ming Yang.

